# EEfinder, a tool for identification of bacterial and viral endogenized elements in eukaryotic genomes

**DOI:** 10.1101/2024.05.24.595709

**Authors:** Yago José Mariz Dias, Filipe Zimmer Dezordi, Gabriel da Luz Wallau

## Abstract

Horizontal transfer is a phenomenon of genetic material transmission between species with no parental relationship. It has been characterized among several major branches of life, including among prokaryotes, viruses and eukaryotes. Genetic elements derived from horizontal transfer are known as Endogenous Bacterial Elements and Endogenous Viral Elements. Endogenous elements characterization provides a snapshot of past host-pathogen interactions and coevolution as well as a reference sequence information to remove false positive results from viral metagenomic studies. However, there is a current lack of standardized tools for endogenous elements identification, which hinder comparative studies and reproducibility within this field. Here we describe EEfinder, a new tool for identification and classification of endogenous elements derived from horizontal transfer. The tool was developed to include six standard steps performed in this type of analysis: data cleaning, pairwise alignment, filtering candidate elements, taxonomy assignment, merging of truncated elements and flanks extraction. We evaluated the sensitivity of EEfinder to identify endogenous elements through comparative analysis using data from the literature and showed that EEfinder can systematically identify endogenous elements with bacterial/viral origin.

## INTRODUCTION

Numerous viruses and bacteria inhabit eukaryotic species, living in close proximity within their host bodies/cells. Virus, Bacteria and Eukaryotes interact in many different ways including at the genomic level through horizontal transfer where fragments or complete genes are swapped between species giving origin to Endogenous Viral Elements (EVEs)^1^ and Endogenous Bacterial Elements (EBEs)^2^. This phenomenon of genetic material transfer between non-parental organisms is called Horizontal Gene Transfer (HGT), which occurs across all major branches of life^3^. Horizontally transferred elements can be vertically inherited if they integrate into germline cells of the host genome^4^. EBEs and EVEs have been increasingly recognized as an important source of evolutionary novelties such as piRNAs produced from EVEs acting as an antiviral defense in mosquitoes and EBE acting as a new sexual chromosome in *Armadillidium vulgare*^2,5^. Moreover, EVEs can be used to calibrate more precisely the evolutionary dating of viral families^6^ owing to the lower mutational rate compared to circulating viruses ^6^ acting as typical viral fossils^1,6,7^.

The endogenization of viral and bacterial genomes into eukaryotic host genomes is currently well-recognized but the exact endogenization mechanisms remain unclear. Transposable elements (TEs), which are abundant in eukaryotic genomes and capable of self-replication and non-homologous recombination, may act as vectors for viral endogenization^8^. Supporting this hypothesis, EVEs endogenization in mosquito genomes is often found in TE-rich regions^5^. TEs may also participate in the generation of virus-TE double-strand RNA and DNA hybrids, facilitating integrase recognition and integration of these chimeric molecules into the eukaryotic genome^9^. Alternatively, viral and bacterial genetic material can integrate into host eukaryotic genomes through non-homologous end joining repair mechanisms^10^.

After integration, the genetic material may mutate and fragment or remain conserved and under purifying selection depending upon the fitness impact on the host^4^. For example, the syncytin gene, which mediates placental cytotrophoblast fusion, shares 100% identity with the Human endogenous retrovirus-W (HERV-W)^11^, syncytin gene was likely integrated into the ancestor of placental mammals around 25 Mya^12^. On the other hand, Leclercq et al. identified a 3 MB insertion of a *Wolbachia* bacteria genome in the *Armadillidium vulgare* host genome, which gave rise to a new sex chromosome^2^. EVEs and EBEs can be transcriptionally active, producing RNA molecules that may be wrongly identified as generated by bona fide independent organisms’ genomes. It can lead to false positives in viral metagenomics and surveillance studies^9,13^. Similarly, endogenous elements from *Wolbachia* genomes can bias studies on this group of bacteria in different eukaryotes^14^.

Precise characterization of EBEs and EVEs is essential to distinguish true infections from endogenous element activity. However, there is no standard method for systematic screening of these elements^15^. An automated approach is needed for research groups to identify and characterize EVEs or EBEs in eukaryotic genomes and differentiate them from actual viral and bacterial infections. To address this, we developed EEfinder, a CLI Python package for systematic identification and characterization of endogenous elements in eukaryotic genomes.

## MATERIALS AND METHODS

### EEfinder Development

EEfinder was developed in Python 3.9 and incorporates six main steps commonly applied in endogenous elements screening^5,9,16,17^. It requires four input files to work (**Figure 1** and **SupplementaryTable1**): a multifasta file of the eukaryotic genome, a multifasta file of virus/bacterial protein database, a metadata table with taxonomic annotation fields (access id, species, genus, family, molecule type, protein product, host), and a host genes baits fasta file with host gene proteins to filter out false positives endogenous elements.

**Figure 1:**
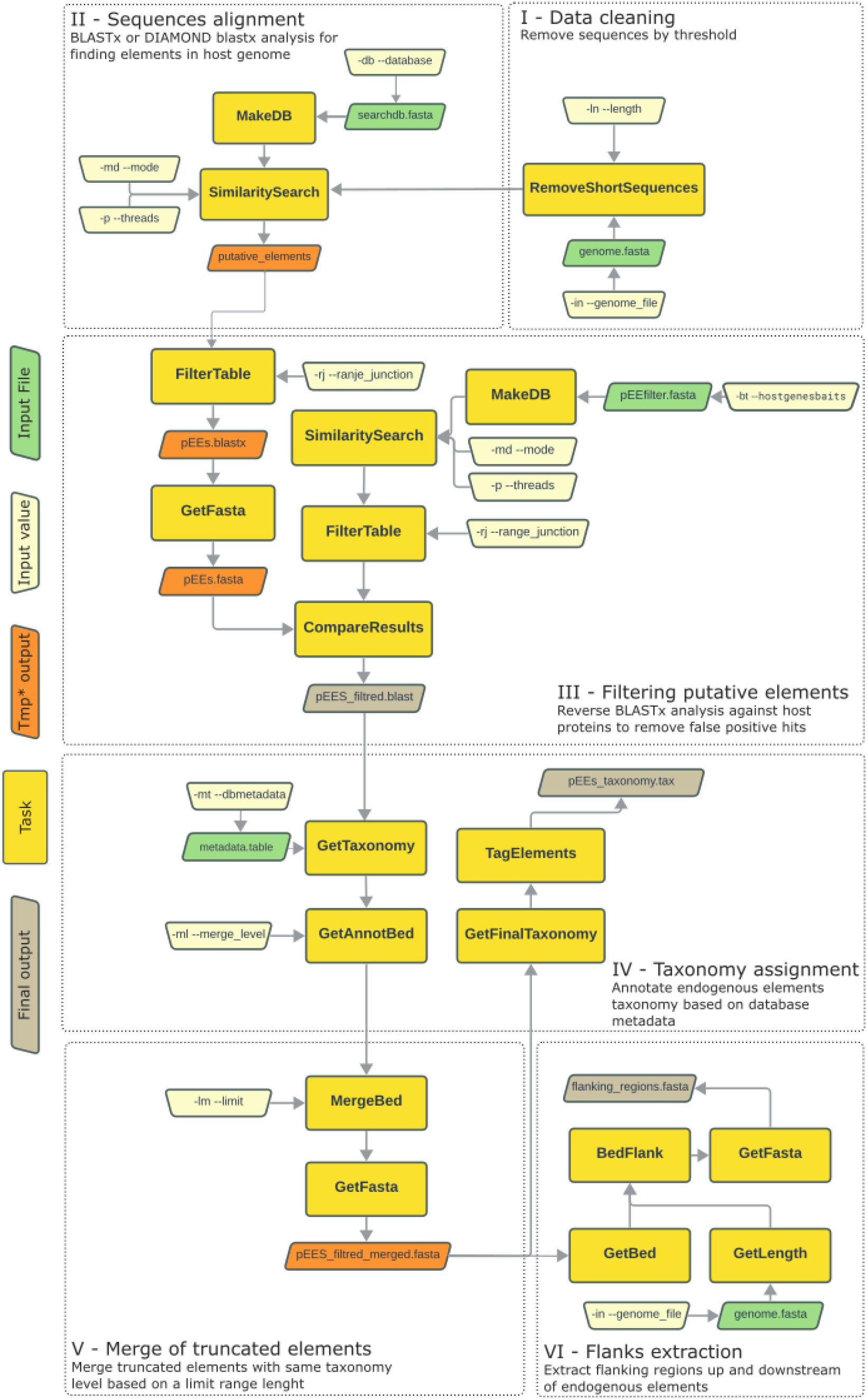
The EEfinder six steps from data input/cleaning to taxonomy assignment and final sequences extraction. *Temporary output files. At default settings, the temporary outputs are deleted, but the user can set to keep those files.

#### Data cleaning

A filter for small genomic sequences was implemented by setting a minimum contig/scaffold length used in the similarity search step (**Figure 1-I**). This approach is used in other studies^5,18^ for filtering out possible contamination and elimination of short contigs that may hinder additional analysis that requires flanking sequences to validate true EEs.

#### Similarity Search

EEfinder was developed with two pairwise-alignment algorithms: BLAST v2.5.0^19^ or DIAMOND 2.0.15^20^, allowing the user to choose between them. We implemented these both software since they have substantial differences in running time and sensitivity to detect EEs. (**Figure 1-II**).

#### Putative Elements Filter

Some bacterial or viral proteins share ancient common ancestors with eukaryotic proteins (e.g., DNA polymerases)^9,17^. Therefore, putative elements identified through pairwise alignment require refinement to filter out false positives using eukaryotic proteins databases. This step is normally performed manually in several published studies^9,17^. EEfinder automates this process by running a reverse alignment against the host protein database, the host genes baits file (**Figure 1-III**). By comparing the initial alignment and the reverse alignment, EEfinder retains only those elements with higher bitscores against the bacterial/viral database than the host proteins database.

#### Taxonomic assignment

In the next step, the EEfinder uses the virus/bacterial metadata table provided by the user to transfer the taxonomic information of each EE based on the most closely related viral/bacterial protein (**Figure 1-IV**). The tool documentation has instructions to build this file and also includes an additional script for automating the table building if the user data is provided.

#### Merging Fragmented Elements

Endogenous elements may be fragmented over evolutionary time by the accumulation of mutations and insertion^4^ or as a result from defective viral particle integration^15^ appearing as degenerated and separated flanking EEs loci. EEfinder merges cis elements identified by the same viral/bacterial family or genus that are apart from a specific range defined by the user (**Figure 1-V**).

#### Flanks Extraction

In the final step, EEfinder extracts the flanks of each element, with the length adjustable by the user (**Figure 1-VI)**. These flanks can be utilized in further analyses, such as gene orthology, transposon searches, or host gene identification^17,21^.

### Alignment Tools Benchmark

To compare the use for each pairwise-alignment mode (BLAST and DIAMOND), we performed tests using different alignment modes: BLASTx and DIAMOND blastx on fast, sensitive, mid-sensitive, more-sensitive, very-sensitive and ultra-sensitive modes. After that, we compared the number of EVEs retrieved, the identity range and the time to run each analysis.

### Computational Resources Usage

To determine the computational resources demand for EEfinder analysis, we ran additional tests with bacterial and virus databases with limited computational resources. We configured four test groups with varying thread counts and corresponding RAM memory: 4 threads with 8 GB, 8 threads with 16 GB, 16 threads with 32 GB, and 32 threads with 64 GB. Each test was conducted three times to estimate the variability, mean and standard deviation.

### Endogenous Viral Elements Validation

EEfinder was benchmarked using the Aag2 genome GCA_021653915 from *Aedes aegypti* (**SupplementaryTable1**). Two viral protein databases were employed: all viral proteins available on NCBI Virus RefSeq (updated September 8, 2022) and proteins described in the study by Whitfield *et al*^17^. This study was chosen once it provides some automated scripts that are widely used in other studies^22,23^. To compare EEfinder results with those from Whitfield *et al*, we updated the taxonomy of EVEs identified by Whitfield et al. using BLASTx analysis against the NCBI Virus RefSeq proteins. The host protein database was constructed by downloading all *Ae. aegypti* proteins available on NCBI RefSeq (updated September 8, 2022). To assess the impact of hypothetical/uncharacterized proteins on the putative EVEs filter, we created two versions of this database: one without any filtering and another excluding all proteins labeled as “hypothetical” or “uncharacterized.” The results comparison were acquired with the Whitfield described proteins database and utilizing the merge level (-ml) set to family.To verify the results differences between Whitfield *et. al*. and EEfinder we used in-house scripts (**SupplementaryTable2**, see **Data Availability** section), that analyzed if the start and end position of each element overlapped with a maximum 100 nt of difference.

In order to check if Whitfield *et al* described fragmented EVEs as separate endogenization events, we extracted the ORFs of each element using ORFfinder^24^ and aligned against the closest viral gene with MAFFT v7^25^, verifying if the ORFs displayed continuity. This process was done for two cases for each family that displayed overlapping elements merged by EEfinder.

### Endogenous Bacterial Elements Sensitivity Validation

To evaluate the applicability of EEfinder for endogenous bacterial elements (EBEs) detection, we aimed to reproduce the results from Leclercq et. al^2^ that described the insertion of a bacteria of the genus *Wolbachia* (wVulC) found integrated in the genome of a pill bug *Armadilium vulgare*. The data used for tests are described in **SupplementaryTable1**. We utilized the WvulC strain as a bacterial database, this strain showed in the Leclerq et al manuscript to be the most similar *Wolbachia* genome to the endogenized one. The following line was used in this analysis:

We tested different parameters for limit merge (-lm) and range junction (-rj). For the range of bases, 10000 and 100 were optimal to recover results similar to the Leclercq study. The merge level (-ml) was set to genus because we used a database with only proteins of the *Wolbachia* genus. After the analysis, we compared the results using the range between flaking EEs as well as the length of the element.

## RESULTS AND DISCUSSION

### EEfinder

The tool is available at (https://github.com/WallauBioinfo/EEfinder), with a Wiki with the full documentation, which includes installation, dependencies versions, testing - with test dataset included in the repository - and documentation about different arguments as well as a short guide to build the input files of the tool. The EEfinder is currently in the first production version (v1.0.0) following the semantic versioning specification^26^. All tests described in the following sections are performed using the EEfinder v1.0.0.

### Alignment tool benchmark

Between the DIAMOND modes, very sensitive mode had the best sensibility, recovering 225 EVEs showing an identity range of 16.2% to 100%. This analysis took 42 hours and 34 minutes to finish. The fast mode retrieved 126 elements, 16.9% - 100% of identity in 8 hours and 6 minutes the best time. Although BLAST showed higher sensitivity with 481 elements with a 10.2% - 100% identity range in 56 hours and 14 minutes (**Figure 2A**).

**Figure 2:**
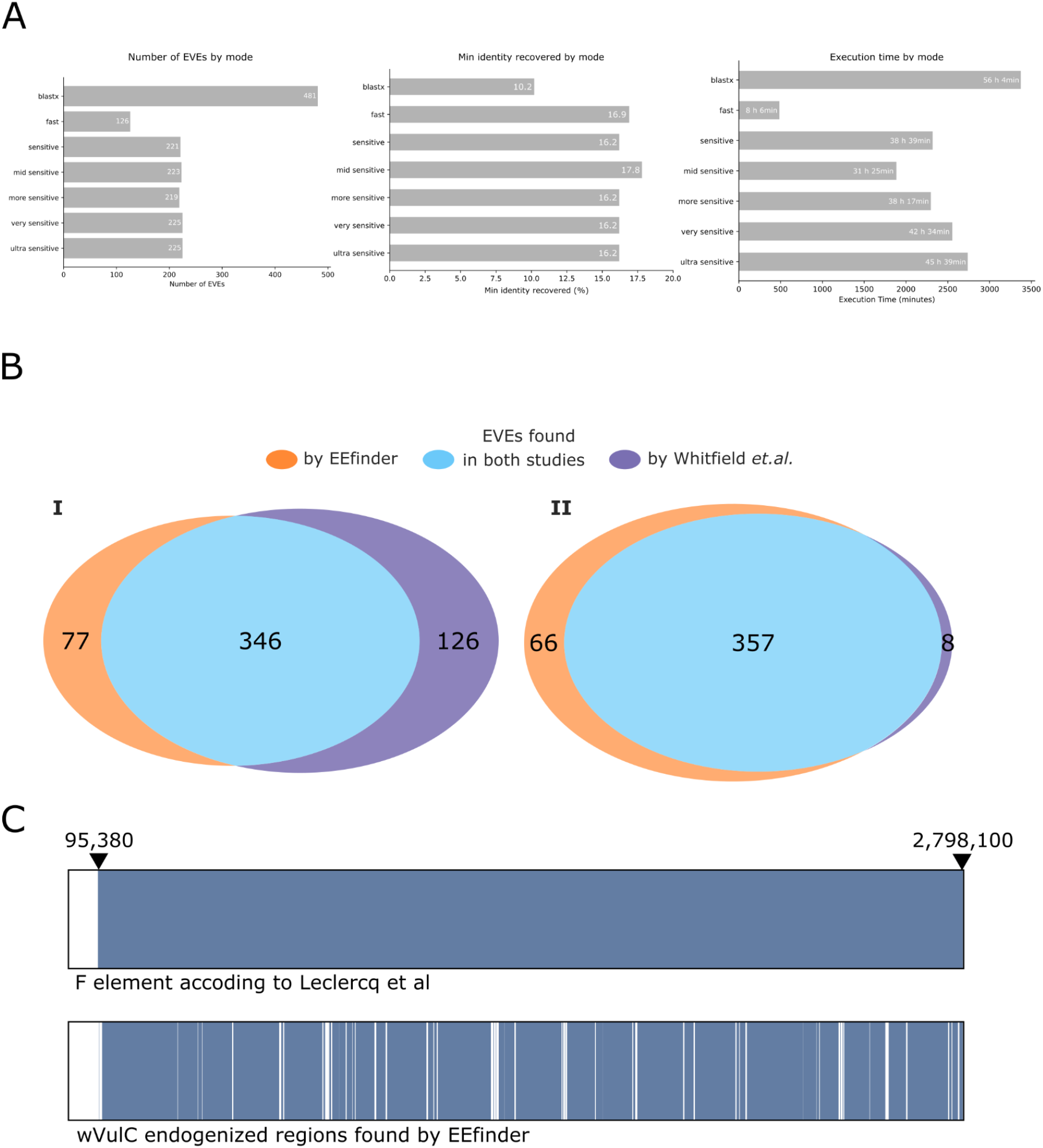
EEfinder benchmark alignment tools and validation. **A**. BLASTx and DIAMOND methods comparrison - number of EVEs recovered using different methods set up. **B**. Comparison between EEfinder and Whitifiled *et. al*.^17^ regarding the number of EVEs recovered by each strategy and the results overlap. **B-I**. Comparison without curation of degenerated elements. **B-II**. Comparison with curation of degenerated elements. **C**. Region of endogenized wVulC identified by Leclercq *et. al*.^2^ and EEfinder on the scaffold 1 of *A. vulgare*. The matched regions found in each study are displayed in blue, the absence of a *Wolbachia* match is represented in white.

In the descriptive study of DIAMOND^20^, the authors stated that the use of a reduced amino acid alphabet doesn’t alter sensitivity, but we showed a clear impact on the pairwise alignment of highly divergent sequences, as is expected in EVEs studies^9^.

### Computational resources usage

The shortest processing time in the resources benchmark test was using 8 threads and 16 GB of memory with a mean of 2 hours and 20 minutes for the virus dataset, meanwhile, the longest time (mean of 2 hours and 38 minutes) was using 4 threads and 8 GB of memory (**Table 1**).

**Table 1:**
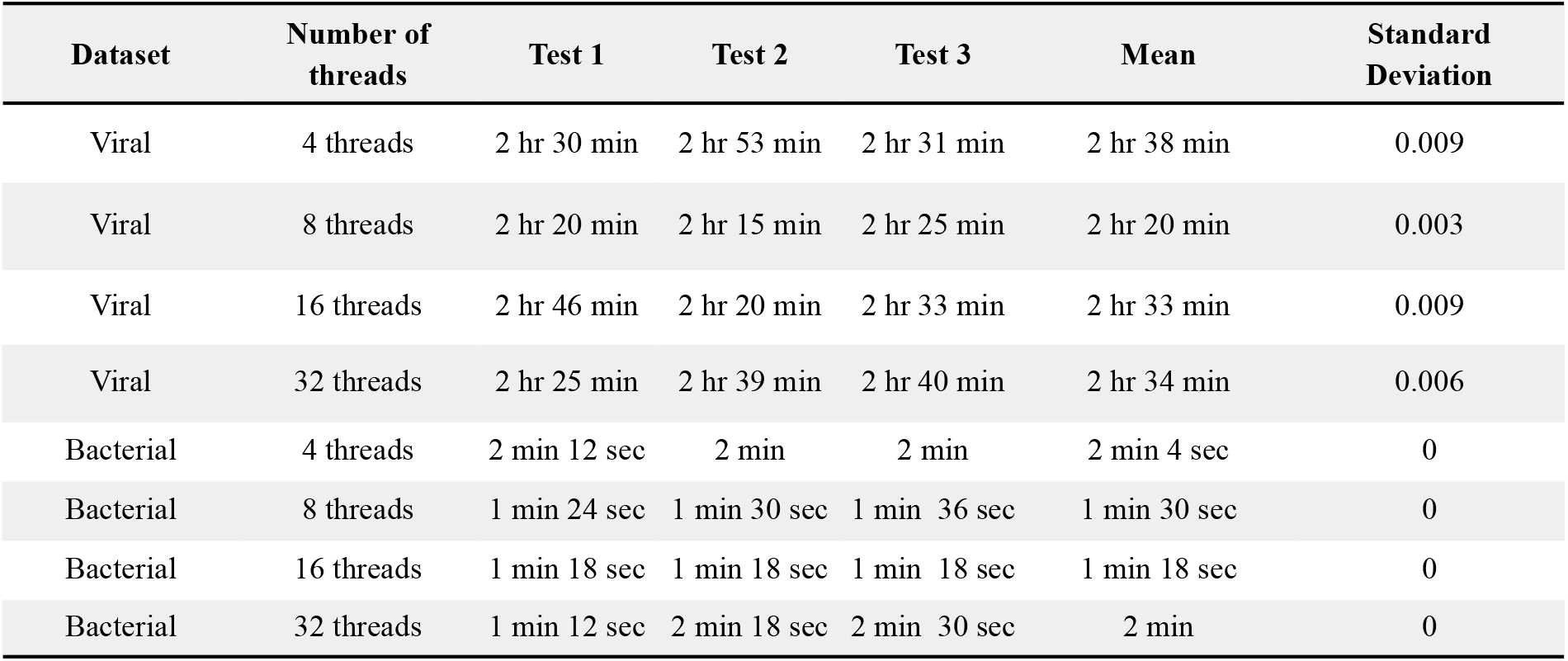
Computational resources comparison of EVEs analysis and EBEs analysis on different resource configurations.

| Dataset | Number of threads | Test 1 | Test 2 | Test 3 | Mean | Standard Deviation |
| --- | --- | --- | --- | --- | --- | --- |
| Viral | 4 threads | 2 hr 30 min | 2 hr 53 min | 2 hr 31 min | 2 hr 38 min | 0.009 |
| Viral | 8 threads | 2 hr 20 min | 2 hr 15 min | 2 hr 25 min | 2 hr 20 min | 0.003 |
| Viral | 16 threads | 2 hr 46 min | 2 hr 20 min | 2 hr 33 min | 2 hr 33 min | 0.009 |
| Viral | 32 threads | 2 hr 25 min | 2 hr 39 min | 2 hr 40 min | 2 hr 34 min | 0.006 |
| Bacterial | 4 threads | 2 min 12 sec | 2 min | 2 min | 2 min 4 sec | 0 |
| Bacterial | 8 threads | 1 min 24 sec | 1 min 30 sec | 1 min 36 sec | 1 min 30 sec | 0 |
| Bacterial | 16 threads | 1 min 18 sec | 1 min 18 sec | 1 min 18 sec | 1 min 18 sec | 0 |
| Bacterial | 32 threads | 1 min 12 sec | 2 min 18 sec | 2 min 30 sec | 2 min | 0 |

For the bacterial dataset, we had the lowest processing time (1 minute and 18 seconds) with 16 threads and 32 GB of RAM. The longest time was with 4 threads and 8 GB of memory, totalizing 2 minutes and 4 seconds (**Table 1**). In conclusion, EEfinder does not require significant computational resources and hence it is suitable to be used even on personal computers.

### EVEs screening

In our first test with the refseq including proteins annotated as “uncharacterized” or “hypothetical” in the host genes baits database against NCBI virus refseq proteins, we encountered 377 EVEs (**SupplementaryTable3**). On the other hand, in the tests with the same host genes baits database filtering uncharacterized and hypothetical proteins, we retrieved 578 elements. When we use only the proteins described by Whitfield *et al* (472 elements)^17^ we were able to characterize 423 elements.

Probably those “uncharacterized” and “hypothetical” proteins were actual unclassified EVEs un, highlighting the necessity to take into consideration the EVEs annotation on eukaryotic genomes. Furthermore, it is important to initiate a review effort for those host protein databases in order to reclassify these proteins that lack annotation.

In order to validate if merging flanking elements with the same taxonomic assignment is biologically meaningful, we extracted the ORFs of both studies and aligned them with MAFFT v7 software^25^ against the best match protein. The validation process was done with 6 merging events (**SupplementaryFigure1**). We were able to show that in three distinct examples (*Blueberry necrotic ring blotch virus, Citrus Leprosis* and *Shayang Fly Virus 1)* Whitfield and collaborators identified a single degenerate element as multiple single endogenization events (**SupplementaryFigure2**). It is noteworthy to point out that EEfinder has merged elements where ORFs were arranged in sequence, one after another, suggesting that these elements were likely derived from a single integration/endogenization event (**SupplementaryFigure2**). This difference results from the merge function of EEfinder, which merges cis elements with the same taxonomic classification within the base pair range defined by the user. Comparing regions found by Whitfield and EEfinder before the curation above mentioned, 343 elements were found in the two analyses, with 24 exclusive elements detected by Whitfield and 80 exclusive elements detected by EEfinder (**Figure 2B-I**). Considering the curated dataset, there are 357 common elements in the two analyses, with 8 exclusive elements detected by Whitfield and 66 exclusive elements detected by EEfinder (**Figure 2B-II**).

EVEs found belong to three most abundant families: Rhabdoviridae, Flaviviridae and Chuviridae. Of the 13 families encountered, 5 were founded only by EEfinder: Spinareoviridae, Virgaviridae, Sedoreoviridae, Closteroviridae and Bromoviridae (**SupplementaryFigure3**). Whitfield had 1 exclusive family (Reoviridae). The number of elements in EEfinder followed the same distribution of Whitfield *et al* analysis (**SupplementaryFigure3**). The proteins retrieved from EEfinder vary in length between 500 to 2000 pb (**SupplementaryFigure4**). The family Kitaviridae showed many elements with length superior to 2000 pb, with an element bigger than 5000 pb.

The total bases retrieved in each study were similar, with a difference of 79.189 bp (**SupplementaryTable4**). The total of bases retrieved by EEfinder was 405,550 pb and Whitfield *et. al*. identified 326,364 bp. The results were consistent between Whitfield and our study regarding the overall EVE content in the Aag2 genome. Moreover, the merging of elements by EEfinder could potentially yield more precise results with improved biological significance.

### EBEs screening

Comparing our results with Leclercq’s study^2^, the test against the isopod *Armadillidium vulgare* genome using the *Wolbachia* wVulC proteins resulted in a single endogenization event in scaffold 1 with the same start and end loci position defined by Leclercq 95,380 - 2,798,100 (**Figure 2C**). Even with the fragmentation displayed by EEfinder, the tool could detect the integration event showing its applicability to identify EBEs in a reproducible way. But the Leclercq *et. al*. study does not clarify if the element found is fragmented as shown by EEfinder. This event may be a result of the accumulation of mutations (SNPs and indels) after the integration event.

## Conclusion

In this study, we developed the first automated tool with a general purpose for the identification of endogenous elements in eukaryotic genomes. The EEfinder sensitivity validation tests have proven the tool’s capability to find endogenous elements replicating results available in the literature for EVEs and EBEs. The tool comes with two algorithms for similarity analysis that can be used for different purposes, BLASTx with the highest sensitivity for identifying all EEs content in one genome. On the other hand, DIAMOND has a lower sensitivity with a lower runtime; these features allow more flexibility to the end user. EEfinder has low computational requirements, it can be used even in low-end personal machines. The tool demonstrates an improvement in the methodology of the EE field using features that enhance reproducibility such as customizable parameters, log files, and fully automated analyses. It is expected that the tool will be widely used in the study of endogenous elements, contributing to the construction of databases for refining surveillance and metagenomic studies, as well as generating data for host-pathogen interactions and pathogen evolution studies.

## Data Availability

A notebook with all lines of code used in this study is available here https://benchling.com/s/etr-2IJ9eTlQjSqtt9qQ4T0G?m=slm-YwVruBwlIl750frj5Pmm. A jupyter notebook with python scripts for data treatment and plots can be found here and the source code of the EEfinder can be found here: https://doi.org/10.6084/m9.figshare.25864525.v1. The branch of EEfinder with the current version of the code can be found here: https://github.com/WallauBioinfo/EEfinder. The NCBI access code for genomes used in this study: *Aedes aegypti* Aag2 GCA_021653915 and *Armadillidium vulgare V1* GCA_001887335.1.

