## Supplementary Tables 1-2-3 and Figures 1-2-3-4 for "EEfinder, a tool for identification of bacterial and viral endogenized elements in eukaryotic genomes"

SupplementaryTable1

| **File** | **Content** | **Usage** |
| --- | --- | --- |
| ae_aegypti_Aag2.fa | Aag2 Genome of *Aedes aegypti (*GCA_021653915) | Similarity Search Benchmark; Resources Benchmark; EVEs Sensitivity Tests |
| ae_aegypti_proteins.fa | Characterized proteins of *Aedes aegypti* (2023-05-30) | Similarity Search Benchmark; Resources Benchmark; EVEs Sensitivity Tests |
| whitfield_protein_database.fasta | Proteins described by Whitfield *et. al.* | Similarity Search Benchmark; Resources Benchmark; EVEs Sensitivity Tests |
| a_vulgare.fa | Genome of *Armadillidium vulgare* (GCA_001887335.1) | Resources Benchmark; EBEs Sensitivity Tests |
| wvulc_proteins.fa | Proteins of WvulC bacteria (2023-05-27) | Resources Benchmark; EBEs Sensitivity Tests |
| a_vulgare_proteins.fa | Characterized proteins of *Armadillidium vulgare* (2023-05-30) | Resources Benchmark; EBEs Sensitivity Tests |

**Supplementary Table 1**: Data used in this study.

SupplementaryTable2

| **Argument** | **Value type** | **Description** |
| --- | --- | --- |
| -in, --genome_file | File | FASTA eukaryotic genome file |
| -db, --database | File | FASTA virus/bacterial protein database |
| -mt, --dbmetadata | File | Table file with data for taxonomic annotation |
| -bt, --hostgenesbaits | File | FASTA host protein file for putative element filter |
| -md, --mode | Text | Selection of which algorithm (BLAST or DIAMOND) will be utilized |
| -ln, --length | Integer | Minimum length of contigs in the genome |
| -fl, --flank | Integer | Desired length for flank extraction |
| -lm, --limit | Integer | Threshold for merging elements with the same taxonomy |
| -ml, --merge_level | Text | Taxonomy level for merging elements |
| -rj, --range_junction | Integer | Junction interval for redundant hits |
| -mp, --mask_per | Integer | Percentage threshold of lowercase characters to consider an EE in a repetitive region |
| -cm, --clean_masked | Flag | Use this argument to remove EEs in repetitive regions |
| -p, --threads | Integer | Threads to be used in the analysis |
| -rm, --removetmp | Flag | Use this argument to remove intermediary files |
| -id, --index_databases | Flag | Flag to index the databases |
| -pr, --prefix | Text | Prefix to analysis files |
| -od, --outdir | Text | Output path |

SuplementaryTable3: Number EVEs identified throughout tests with different host genes baits databases

| **Viral protein database** | **Host genes baits database** | **Number of EVEs** |
| --- | --- | --- |
| NCBI virus refseq | Uncharacterized host protein database | 377 |
| NCBI virus refseq | Characterized host protein database | 578 |
| Whitfield *et. al.* | Host protein database used by Whitifield et al. | 423 |

Supplementary Table 4:

| **Study** | **Total bases founded** |
| --- | --- |
| EEfinder | 405,550 bp |
| Whitfield et. al. 2019 | 326,364 bp |

**Supplementary Table 4:** Total bases found in each study.

SupplementaryFigures1

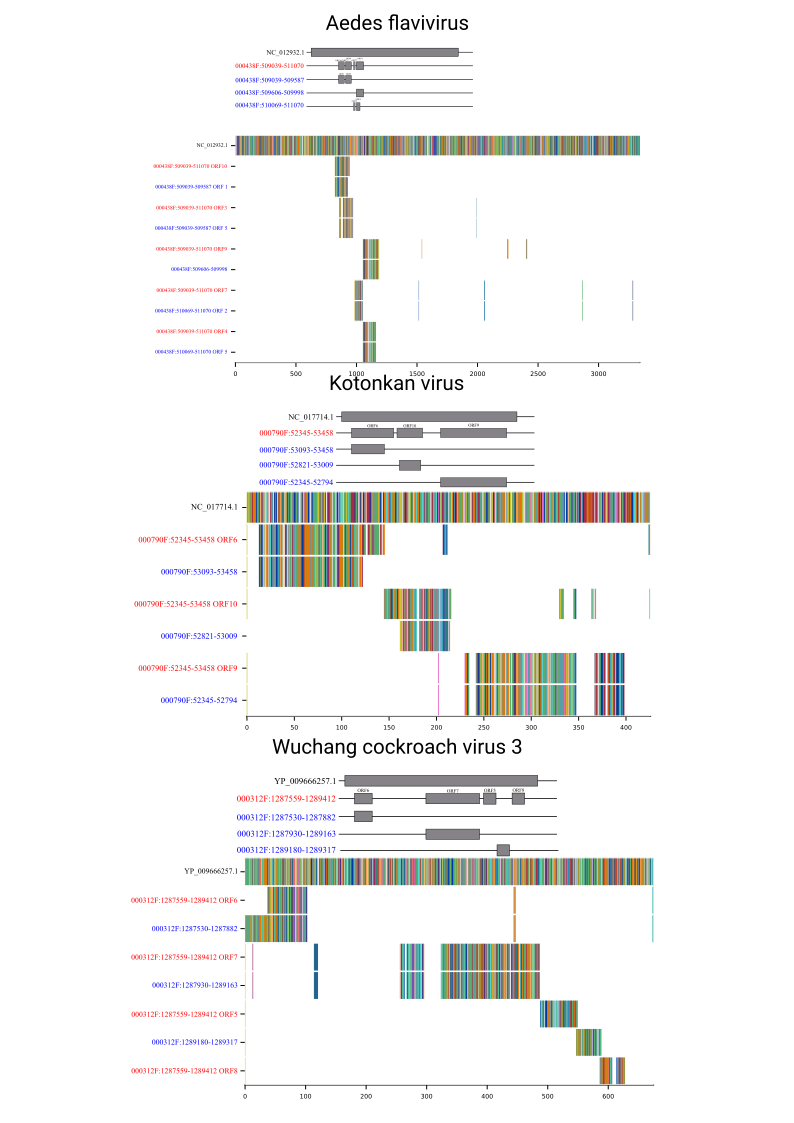

SupplementaryFigures2

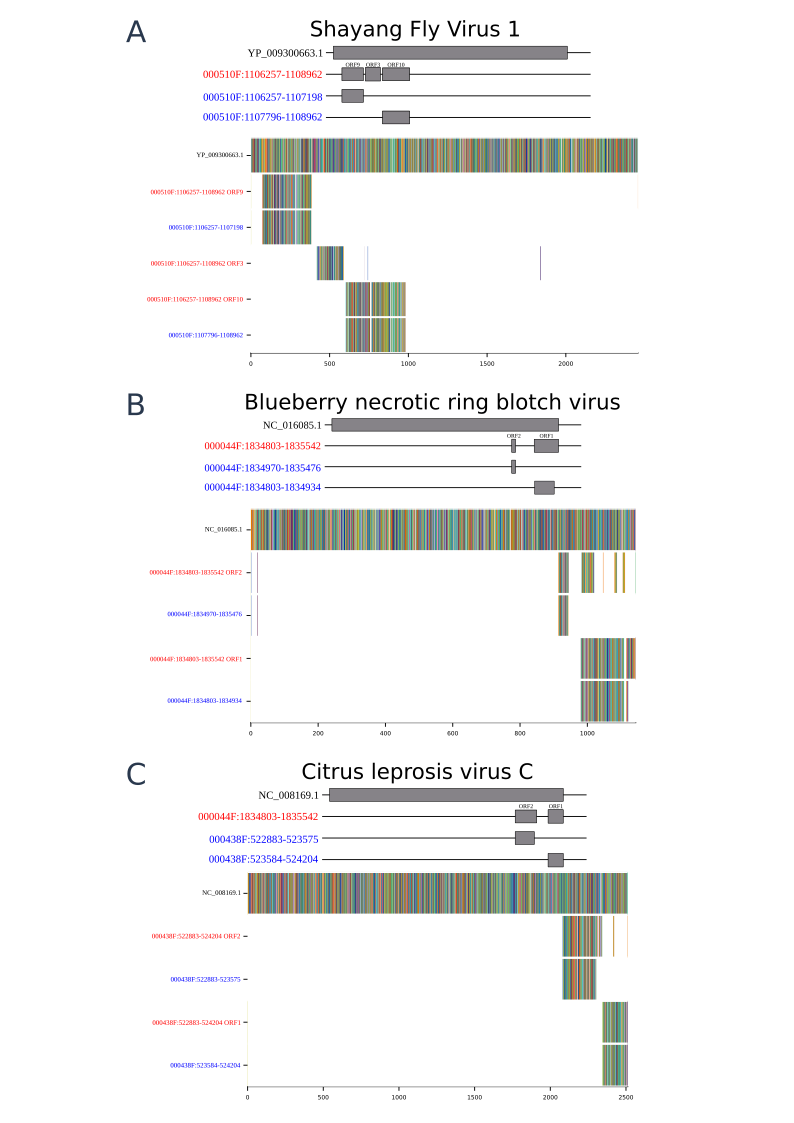

**Figure 2:** Alignments of splitted elements identified by Whitfield et al in blue font, and elements identified by EEfinder in red font. Above each alignment has a representation of the elements found by EEfinder in red font and elements found by Whitfield showing the continuity of ORF’s elements. A) Shayang Fly virus 1 aligned with corresponding EVEs. B) Blueberry necrotic ring blotch virus aligned with corresponding EVEs. C) Citrus leprosy virus C aligned with corresponding EVEs.

Supplementary Figure 3:

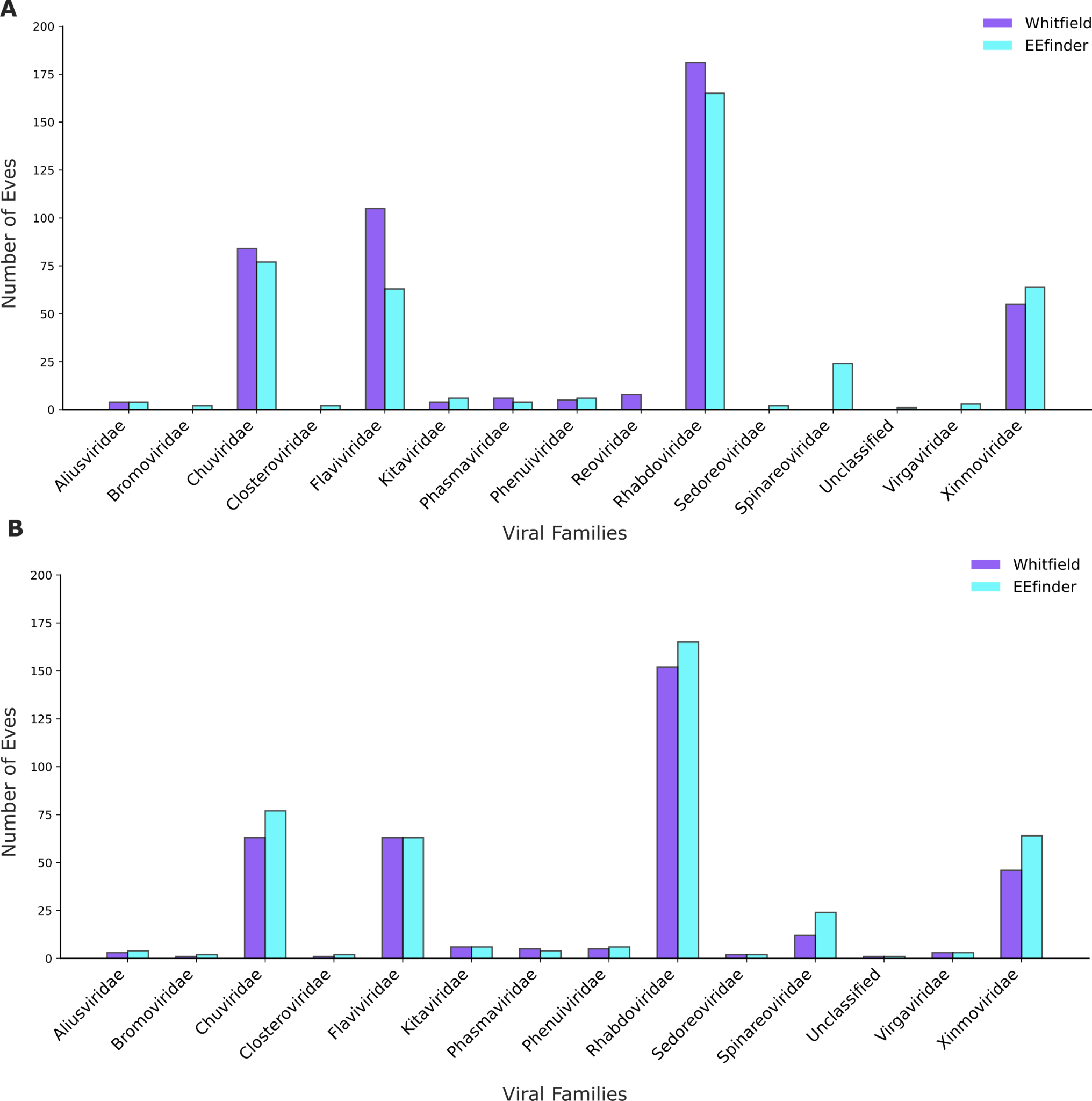

**Supplementary Figure 3**: Comparison between EVEs by family found by EEfinder and Whitifiled EVEs. **A**. Comparison without curation of degenerated elements. **B**. Comparison with curation of degenerated elements.

Supplementary Figure 4:

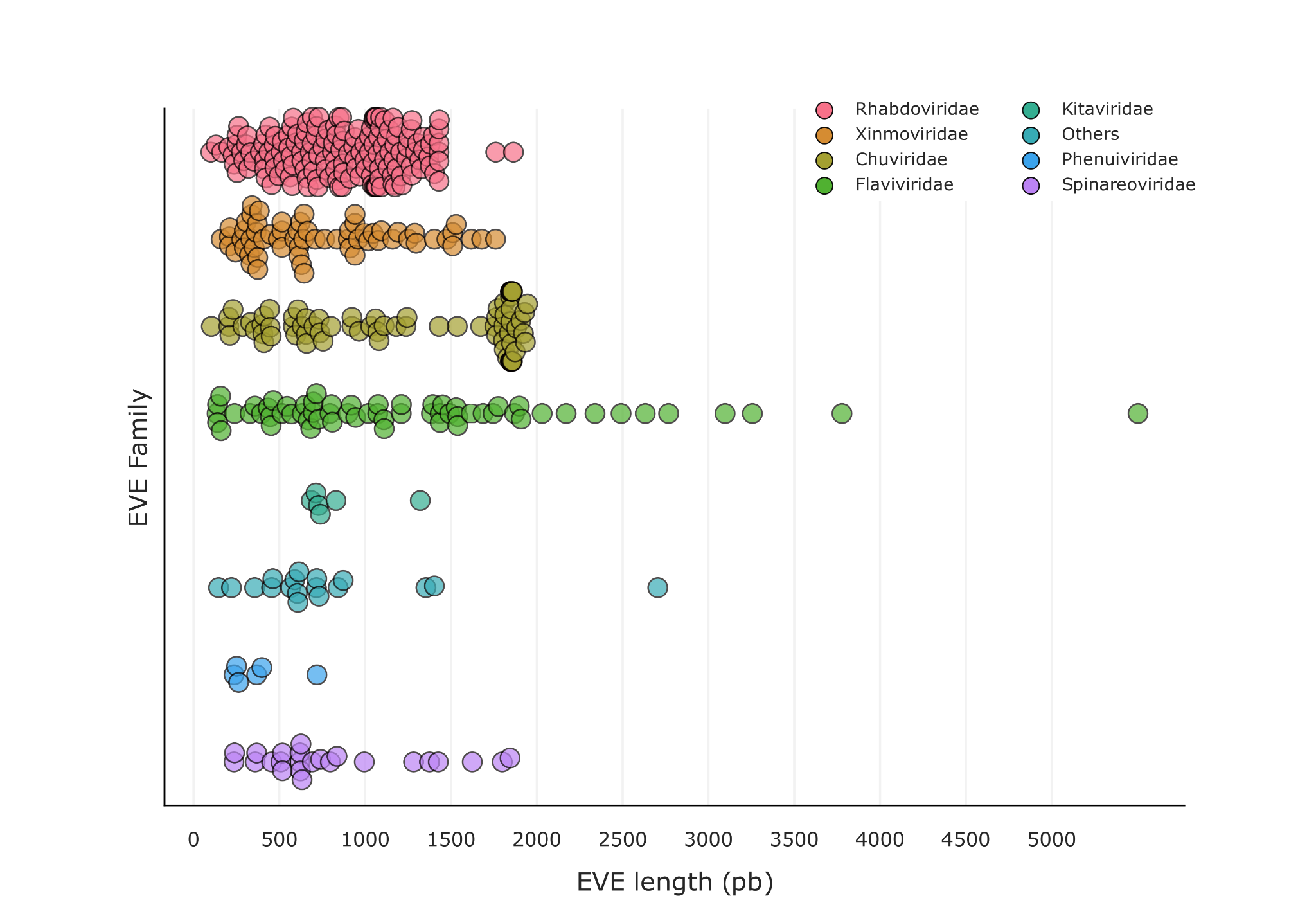

**Supplementary Figure 4:** Display of EVE length divided by virus family.
